# ABISS: An Open-Source, Low-Cost Platform for Auditory and Visual Intrinsic Optical Signal Imaging

**DOI:** 10.64898/2026.08.03.741387

**Authors:** Zhili Qu, Kimia Kazemi, Tianyang Wu, Shambulingayya N. Doddapujar, Tyler A. Marrazzo, Mattia Gazzola, Howard J. Gritton

**Author notes:** Correspondence: Howard J. Gritton. First author (Zhili Qu).

## Abstract

Defining the boundaries of functional cortical areas is increasingly important for targeted electrophysiology, optical imaging, viral delivery, and circuit manipulation. Intrinsic optical signal imaging (IOSI) provides a rapid and minimally invasive approach for mapping stimulus-evoked cortical activity, but its implementation often requires laboratory-specific combinations of stimulus-generation hardware, experiment-control software, synchronization devices, and data-acquisition systems. These requirements limit accessibility and hinder the use of IOSI as a routine functional mapping tool. Here, we present the Arduino-Based Intrinsic Stimulation System (ABISS), an open-source platform that integrates auditory and visual stimulus generation, trial timing, and image-acquisition triggering into a single programmable device. ABISS generates auditory tone stimuli, VGA-based visual stimuli, and tightly synchronized camera-trigger pulses without requiring a dedicated experiment-control system. Stimulus protocols are also fully modifiable in firmware. Performance was evaluated in auditory and visual cortices of mice. Engineering validation demonstrated accurate stimulus generation and synchronization between stimulus delivery and camera triggering over extended recording sessions. Biological validation showed that ABISS output results in auditory and visual intrinsic signal maps comparable to those obtained using highly specialized or commercial platforms. Together, these findings demonstrate the utility of a low-cost open-source platform for experimental control of intrinsic optical signal imaging. By reducing the technical and financial barriers associated with routine intrinsic optical imaging, ABISS facilitates broader adoption of functional cortical mapping as a tool for improved cortical localization in neuroscience experiments.

**Significance Statement:** Functional cortical mapping is an increasingly important element of neuroscience experimental design as anatomical coordinates alone are often insufficient for defining cortical boundaries in individual animals. Intrinsic optical signal imaging provides an effective solution but traditionally requires specialized hardware, commercial stimulus-generation systems, and laboratory-specific synchronization workflows. We developed ABISS, an inexpensive, open-source platform that integrates auditory and visual stimulus generation with synchronized camera triggering in a single programmable device. ABISS produces functional cortical maps comparable to those obtainable with commercial or specialized systems while substantially reducing hardware complexity and cost. By making intrinsic optical signal imaging more accessible, ABISS lowers the practical barriers for routine functional mapping of the brain and promotes adoption of this important neuroscience technique.

## Introduction

Precise identification of functional cortical boundaries is fundamental to neuroscience experiments that require spatially targeted access to specific cortical regions. Traditionally, researchers have used stereotaxic coordinates and brain atlases to guide experimental targeting. Although these atlases provide valuable anatomical references, they are generated from averaged populations and thus do not inherently capture the breadth of inter-animal variability in cortical organization. Indeed, recent studies have shown that anatomical stereotaxic coordinates are often insufficient for precise cortical targeting ^1–3^. As genetic, optical, and electrophysiological tools increasingly enable interrogation of smaller and more specialized domains, the precision required for experimental targeting has increased proportionately. Functional mapping of individual animals prior to targeted experimentation is therefore becoming an increasingly important component for experiments that seek to relate neuronal activity, connectivity, and behavior to specific cortical regions.

Intrinsic optical signal imaging (IOSI) is particularly well suited for this purpose because it provides rapid and minimally invasive functional maps, with relatively simple optical instrumentation. IOSI measures stimulus-evoked changes in tissue reflectance arising from activity-dependent changes in blood oxygenation, blood volume, and light scattering. IOSI can produce functional maps of the cortical surface through the skull without exogenous dyes or genetically expressed indicators^4–6^. Over the past four decades, IOSI has been widely used to investigate the functional organization of sensory systems, including retinotopic, somatotopic, olfactory and tonotopic mapping ^4,7–12^. Because IOSI can rapidly localize functional cortical boundaries across large regions of cortex, it has become an effective method for identifying experimental targets before subsequent electrophysiological recording, viral delivery, pharmacological manipulation, or lesion experiments ^13–17^.

Despite its utility, IOSI is not a routine technique in many neuroscience laboratories. Although the optical requirements are conceptually straightforward, successful implementation typically requires tight coordination among stimulus-generation hardware, experiment-control software, and image acquisition. Consequently, IOSI implementation often relies on laboratory-specific combinations of commercial systems, custom electronics, MATLAB or LabVIEW software, and data-acquisition hardware, that are managed in customized experimental workflows ^13,14,18,19^. Although these configurations are powerful and flexible, distributing experimental elements across multiple platforms increases setup complexity, complicates reproducibility across lab groups, and creates practical barriers for laboratories seeking to use IOSI for routine functional mapping.

Recent open-source efforts have substantially improved the accessibility of IOSI by reducing the cost of illumination, optics, imaging systems, and data analysis ^13,18,19^. However, stimulus generation, trial control, camera triggering, and synchronization across platforms are still commonly distributed across multiple computers and laboratory-specific hardware devices. As a result, implementing even relatively simple IOSI paradigms often requires users to assemble and validate multiple independent hardware and software components before experiments can begin. A compact, reproducible platform that integrates experimental control functions would substantially reduce the technical barriers of routine functional cortical mapping across laboratories.

Here, we present the Arduino-Based Intrinsic Stimulation System (ABISS), a low-cost, open-source platform for IOSI that integrates auditory and visual stimulus generation, experimental timing, and synchronized camera triggering into a single compact programmable device. ABISS uses an Arduino-controlled hardware platform to consolidate the distributed stimulus-control and synchronization infrastructure that accompanies many existing IOSI workflows. Experimental protocols are fully customizable in firmware and delivered through a simple and easy to assemble hardware platform, allowing investigators to adapt stimulus parameters without developing a separate experiment-control interface. We evaluated ABISS at both the engineering and biological levels by quantifying stimulus control and synchronization, comparing auditory cortical maps obtained with ABISS to those obtained using a conventional commercial system for auditory control, and demonstrating visual cortex mapping using the same platform. Together, these experiments establish ABISS as an intuitive, simple, and highly flexible infrastructure for laboratories seeking to implement routine functional cortical mapping without relying on expensive or highly specialized experiment-control systems.

## Materials and Methods

### Hardware Design and Assembly

ABISS consists of a custom printed circuit board, an Arduino microcontroller, and off-the-shelf electrical components that, when assembled, allow integrated stimulus generation, imaging acquisition triggering, and experimental flow control for intrinsic optical signal imaging (IOSI) via a single hardware platform (Figure 1). The system is designed around an Arduino Nano microcontroller that coordinates sensory stimulus delivery and synchronized camera triggering while interfacing with external components (i.e., speakers, VGA monitor, and cameras) through dedicated digital input and output connections. The assembled board is housed within a custom enclosure that protects the electronics while maintaining access to all external connectors (Extended Data Figure 1-1). Power is supplied through a barrel jack by a regulated +5 V DC source. The printed circuit board (Extended Data Figure 1-2) design provides three digital input/output BNC-style coaxial connectors. In the configuration demonstrated here, these connectors provide one input for user- or externally initiated trial triggering, one digital output for camera triggering, and one general-purpose input/output connector reserved for future applications. In the PCB layout, Arduino digital pins D8, D7, and D9 are routed to coaxial connectors J2, J3, and J4, respectively. For the experiments described here, the J4/D9 served as the external TTL start input, J2/D8 served as the synchronized camera-trigger output, and J3/D7 was reserved as a programmable general-purpose I/O line. The connector-to-pin routing is fixed by the PCB, whereas the input/output behavior of each digital line can be modified in firmware. Visual stimuli are generated through an integrated VGA interface directly driven by the Arduino. The device incorporates a MAX660 charge-pump voltage inverter that generates an approximately -5 V rail from the +5 V input, providing the negative supply used by the LT1970 amplifier stage. Auditory stimuli are generated using an AD9833-based waveform-generation module (GY-9833) controlled via a serial peripheral interface (SPI). Signal amplitude is managed by a PT2258 digital volume controller communicating over I^2^C, which allows independent control of precise per-tone amplitude with smooth programmable onset and offset ramps. The resulting signal is amplified by an LT1970 amplifier stage (TSSOP-20) which provides the voltage and current drive required by the external speaker. The assembled hardware and system wiring are shown in Figures 1a and 1b. Complete PCB layouts, circuit schematics, component lists, assembly documentation, and hardware costs are provided in Extended Data Table 1 and Extended Data Figures 1-2 and 1-3.

**Figure 1.**
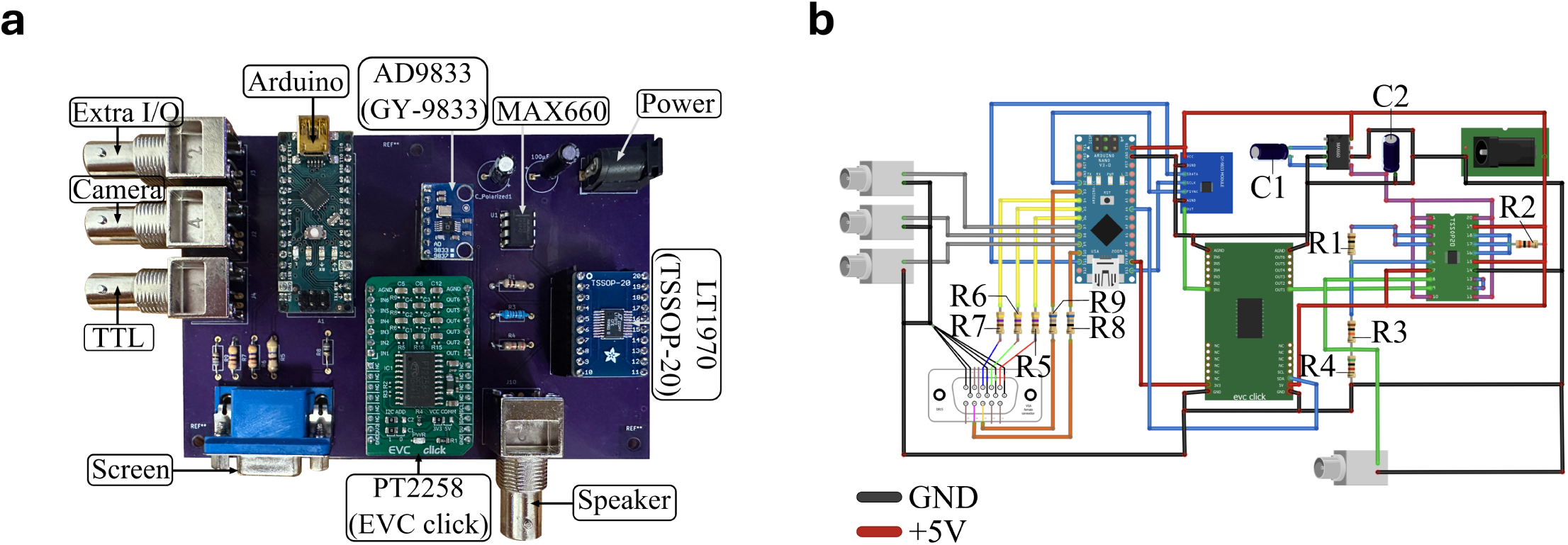
Custom electronic control system for intrinsic optical signal imaging (IOSI). **a.** Photograph of the ABISS board showing the physical layout and component assembly. An Arduino Nano serves as the central microcontroller, interfacing with BNC-style coaxial signal input/output, analog audio output, and a VGA output for visual stimuli. A MAX660 charge pump generates an approximately -5 V supply rail for the LT1970 amplifier. Auditory signals are generated by an AD9833-based DDS module (GY-9833), controlled in amplitude by a digital volume controller (PT2258), amplified by an LT1970 stage (TSSOP-20), and routed to a coaxial output. **b.** Wiring diagram showing the physical implementation of the system, with resistor and capacitor locations labelled. Color-coded wiring indicates signal routing and power connections: crimson red, +5 V power; purple, −5 V power generated by the MAX660 charge pump; black, ground; sky blue, general digital signals; gray, digital I/O connected to the three BNC ports; yellow, VGA video signals (red, green, and blue); orange, VGA horizontal and vertical synchronization signals; and emerald green, analog audio signals. Component values are as follows: R1, 1 Ω; R2, 3 kΩ; R3, 10 kΩ; R4, 5 kΩ; R5–R7, 470 Ω; R8–R9, 68 Ω; C1–C2, 100 *µ*F.

### Experimental Hardware Configuration

ABISS provides independent auditory and visual stimulus-generation. The auditory configuration generates user-defined pure-tone sequences with programmable frequency, amplitude, stimulus duration, and inter-trial timing. The visual configuration produces VGA-compatible output (i.e., high-contrast patterns, including the drifting checkerboard bars used for retinotopic mapping shown in Figure 2c). For standard IOSI experiments, ABISS was configured as illustrated in Figure 2a. An external user-initiated start signal (button press delivered to the ABISS trigger input) initiated the stimulus sequence. Depending on the experiment, the audio output was connected to an external speaker, or alternatively, the VGA output was connected directly to a display positioned in front of the animal. Camera synchronization was provided through the dedicated trigger output of ABISS to a FLIR camera via a general-purpose input/output (GPIO) connector. Camera acquisition was configured in SpinView (Spinnaker SDK, Teledyne FLIR) by enabling trigger mode with rising-edge detection so that each TTL pulse initiated the acquisition of a single image frame. Acquired images were streamed to a desktop computer for storage in real-time.

**Figure 2.**
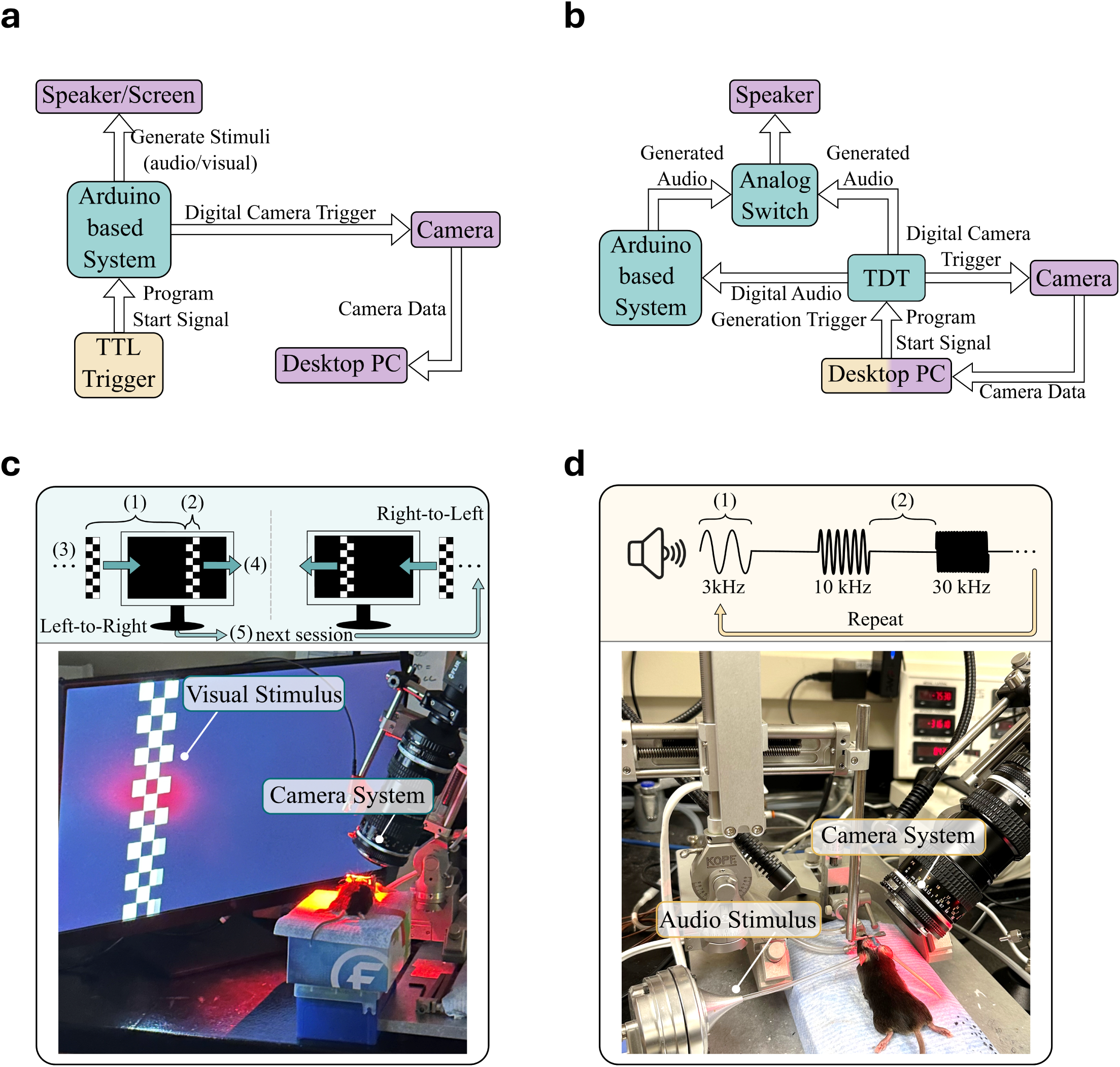
Experimental hardware configurations and sensory stimulation paradigms. **a.** Hardware system setup for IOSI auditory and visual experiments. A user controlled TTL trigger (button press) initiates stimulus presentation, with ABISS generating audio and/or visual stimuli while simultaneously issuing digital pulses to drive camera acquisition. Captured camera data is transmitted to a desktop PC for storage and post-hoc analysis. **b.** Design for integrating ABISS with the TDT RZ6 for tonotopic map comparisons. The RZ6 receives a program start signal from the desktop PC. The RZ6 controlled all elements of the study alternating between generating audio directly and triggering ABISS to generate audio via TTL pulses. Output audio from the two systems was routed through an analog switch box while camera acquisition was digitally triggered by the RZ6 system and recorded on the desktop PC. **c.** Visual stimulation paradigm and experiment. Top: Example visual stimulus generated by ABISS during visual IOSI. The stimulus parameters are programmable in the Arduino firmware, including: (1) inter-bar spacing, (2) bar width, (3) number of bars, (4) bar movement speed, and (5) inter-sweep interval between forward and backward moving-bar presentations. The visual pattern displayed within each bar can also be modified through the firmware. Bottom: Annotated photograph of the visual IOSI experiment. During imaging, the visual stimulus moved across the display from one edge of the screen to the opposite edge while the anesthetized animal was passively exposed to the stimulus. Images were acquired simultaneously at 20 Hz under continuous 625-nm red LED illumination. **d.** Auditory stimulation paradigm and experiment. Top: Example auditory stimulus generated by ABISS during auditory IOSI. Auditory stimulus parameters are programmable in the Arduino firmware, including: (1) tone frequency, sound duration, and (2) intertrial interval. In addition, the auditory output amplitude, onset/offset ramp can be adjusted in Arduino firmware. Bottom: Annotated photograph of the auditory IOSI experiment. The auditory stimulus was generated by ABISS and delivered to an MF1 Multi-Field Magnetic Speaker in a closed-field configuration. A guide tube connected the speaker output to the anesthetized animal’s ear. Images were acquired simultaneously at 20 Hz under continuous 625-nm red LED illumination from the contralateral auditory cortex.

To compare ABISS auditory stimulus generation accuracy, we compared ABISS results to those produced with a commercial system. Auditory imaging experiments were interleaved with a Tucker-Davis Technologies (TDT) RZ6 processor controlled by Synapse software (Figure 2b). Before each experiment, both systems were calibrated to deliver 75 dB SPL pure tones at the animal’s ear. RZ6 and ABISS trials alternated within the same imaging session while the speaker, acoustic delivery path, camera-triggering pathway, imaging setup, and animal preparation remained unchanged. During RZ6 trials, Synapse generated the auditory stimulus directly through the RZ6 hardware. During ABISS trials, Synapse provided a TTL pulse to the ABISS digital trigger input, initiating one programmed tone presentation. Tone frequency advanced sequentially across trigger pulses so that both systems presented matched but independently generated auditory stimuli at 3, 10, and 30 kHz. The two audio outputs were routed through a two-way switch box (DB502-0002, Cable Leader), allowing only the selected system to drive the speaker on each trial. Camera acquisition was triggered by the TDT RZ6 system for both ABISS and RZ6 trials, ensuring identical image-acquisition timing across the two conditions. This interleaved design enabled direct comparison of the cortical response maps evoked by the two systems under otherwise matched experimental conditions.

Because the TDT RZ6 system used for the auditory comparison did not provide an equivalent VGA-based visual stimulus output, a matched side-by-side comparison was not available for the visual experiments. Instead, visual stimulus generation was evaluated by determining whether ABISS produced retinotopic and visual field-sign maps consistent with previously published IOSI studies.

### Subjects

All animal procedures were approved by the University of Illinois at Urbana-Champaign Institutional Animal Care and Use Committee (IACUC) under protocols 23260 and 24059 . Three VIP-Cre transgenic mice (B6J.Cg-*Vip^tm1(cre)Zjh^* /AreckJ; Jackson Laboratory; two males and one female) between 8 and 12 weeks of age were used in this study. One animal was used for the direct comparison of maps between ABISS and the TDT RZ6 auditory stimulation system, one animal was used for the RZ6 self-consistency benchmark testing, and one animal was used for visual intrinsic optical signal imaging experiments.

### Surgery

Mice underwent surgical implantation of a custom head-plate immediately before intrinsic optical signal imaging. Animals remained under anesthesia throughout the imaging experiment. Animals were initially anesthetized with 2% isoflurane and secured in a stereotaxic frame. Following a midline scalp incision, the skull was exposed and bregma was identified as the stereotaxic reference point. A custom head plate was positioned anterior to bregma and secured to the skull using dental cement.

Stereotaxic reference marks were placed on the skull to guide positioning of the imaging field. For auditory imaging, marks were placed approximately 2.5 mm lateral and -2.4 mm and -3.7 mm posterior to bregma. For visual imaging, reference marks were placed approximately 0.75 mm lateral and -3.2 and -4.8 posterior to bregma. These landmarks were used to position the imaging system over the appropriate cortical region before functional mapping.

After the dental cement had cured, the stereotaxic apparatus and anesthesia nose cone were repositioned to provide unobstructed optical access to the cranial surface while maintaining stable head fixation.

### Intrinsic optical signal imaging

Directly before imaging, chlorprothixene (1.5 mg/kg, intraperitoneal) was administered and isoflurane anesthesia was reduced to 0.75%. Images were acquired transcranially through the intact skull, which was maintained optically transparent with sterile saline. Intrinsic optical signal imaging was performed using a tandem-lens macroscope (Nikkor 55 mm 1:2.8 and 85 mm 1:2) coupled to a 16-bit CMOS monochrome camera (BFLY-U3-23S6M-C, Teledyne FLIR). The intrinsic signals were captured at a frequency of 20 Hz under red light illumination (625-nm).

### Auditory Intrinsic optical signal imaging

An auditory IOSI trial consisted of a 1.5 s baseline, followed by a 1 s pure tone stimulus (75 dB SPL) at 3, 10, or 30 kHz ^3^, and a 28.5 s inter-trial interval (Figure 2d). Sound pressure levels were calibrated using an ACO model 4016 microphone, an ACOustical Interface PS9200, and a customized RPvdsEx circuit before each experiment. In the self-consistency experiments (described below), auditory stimuli were generated using the TDT RZ6 processor driving a TDT Multi-Field Magnetic (MF1) speaker in a closed-field configuration. For ABISS comparison experiments, stimulus presentation alternated between the RZ6 system and ABISS while maintaining identical acoustic hardware, speaker placement, and imaging conditions. For each frequency, the two systems alternated stimulus presentation before advancing to the next frequency, allowing direct comparison of evoked intrinsic signal maps under matched experimental conditions.

For the direct ABISS–RZ6 comparison, 10 trials were acquired for each system at each stimulus frequency. In a separate RZ6-only experiment, 20 trials were acquired per frequency to assess self-consistency; these trials were later divided into odd- and even-trial subsets containing 10 trials each.

### Visual Intrinsic optical signal imaging

Visual cortical mapping was performed using phase-encoded drifting checkerboard bar stimuli presented on a VGA display (440 × 480 pixels) positioned 15 cm from the contralateral eye and rotated approximately 20° inward to align with the natural visual axis of the mouse, consistent with previous mouse IOSI protocols^13,18^. Vertical and horizontal checkerboard bars swept across the display in opposite directions to generate retinotopic maps of azimuth and elevation.

Stimuli consisted of 10 drifting checkerboard bars presented on a black background (Figure 2c). Vertical bars moved horizontally across the display to map azimuth, whereas horizontal bars moved vertically to map elevation. Bars were updated at 20 Hz to produce smooth horizontal or vertical sweeps across the display. The bars moved at approximately 25 pixels/s and required approximately 19.2 s to traverse the 480-pixel display field. A unidirectional sweep was defined as the complete traversal of the 10-bar train across the display in one direction. After the forward sweep, the display was blanked for 20 s, after which the bar train moved in the opposite direction. The forward and reverse sweeps together constituted one bidirectional sweep cycle. Nine bidirectional sweep cycles were acquired for each stimulus orientation and used for subsequent retinotopic and visual field-sign analyses. Camera-trigger pulses were generated at 20 Hz throughout each active unidirectional sweep and were paused during the 20 s inter-sweep blank interval. Because imaging was performed over the right hemisphere, animals were positioned so that the left eye faced the center of the display, with the optic axis approximately perpendicular to the screen.

### Engineering Validation

In addition to assessing the consistency of biological maps relative to commercial or previously published systems, we also characterized the temporal and spectral performance of ABISS hardware synchronization via engineering validation experiments. Camera trigger outputs and audio or visual stimulus signals were recorded simultaneously using a TDT RZ6 system running a customized Synapse acquisition circuit sampled at 97.6 kHz. Camera trigger pulses were acquired through a digital input channel, while the audio outputs were recorded through an analog input channel. For visual validation, the VGA red-channel output (pin 1) and ground (pin 6) were routed to the RZ6 via a custom adapter to provide an analog representation of stimulus timing. For auditory validation, the trigger protocol was modified to include a 2 s pre-stimulus period, a 2 s tone presentation, and 2 s post-stimulus period to facilitate temporal alignment measurements. Validation recordings were analyzed offline to quantify stimulus spectral content and synchronization between stimulus delivery and camera triggering (Figure 3).

**Figure 3.**
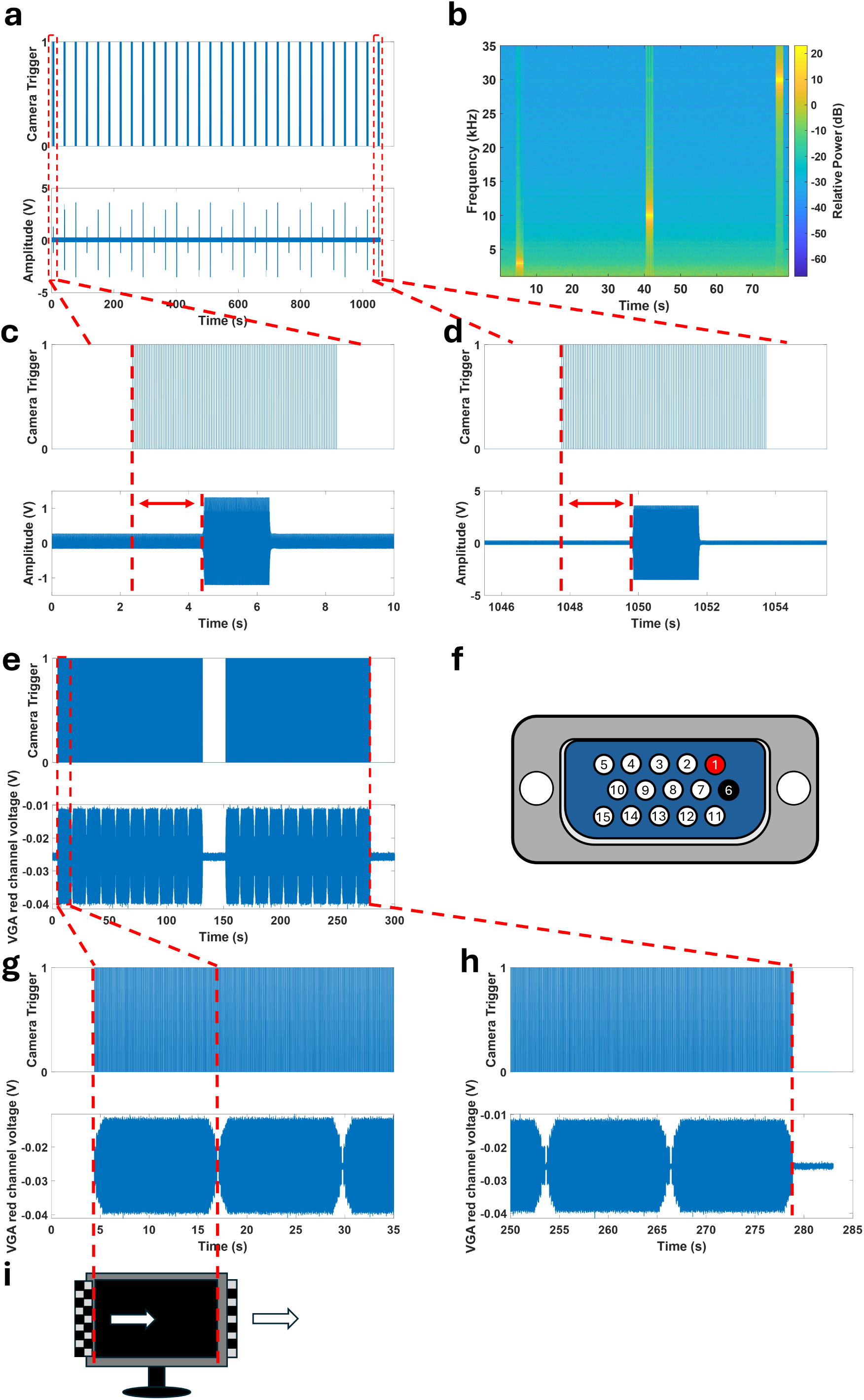
Characterization of ABISS-generated auditory/visual stimuli and camera trigger synchronization. **a.** Example auditory stimulus sequence from a typical imaging session. Tones at 3, 10, and 30 kHz were each calibrated to 75 dB SPL and presented sequentially, with voltage amplitude adjusted per frequency to account for speaker response properties. The three-frequency sequence was repeated 10 times for a total of 30 tone presentations. For visualization purposes, the camera acquisition trigger was set to begin 2 s before tone onset and end 2 s after tone offset, with a tone duration of 2 s. **b.** Spectrogram (squared magnitude of the short-time Fourier transform, STFT) of the recorded tone sequence, confirming accurate frequency content across all three designed tones. **c.** Camera trigger timing associated with the first 3 kHz tone presentation. The measured pre-tone trigger interval (red double arrow) was 2.00045 s, closely matching the intended 2-s delay. **d.** Camera trigger timing associated with the final 30 kHz tone presentation, occurring after approximately 1000 s of continuous operation. The measured pre-tone trigger interval (red double arrow) was 2.09 s, reflecting a cumulative timing drift of approximately 0.09 s relative to the theoretical value — less than 0.01% of the total session duration — demonstrating highly reliable synchronization between audio stimulus delivery and camera acquisition throughout the session. **e.** Example visual validation recording showing one bidirectional sweep cycle. During the forward unidirectional sweep, a train of 10 checkerboard-patterned bars traversed the display from left to right. Because the bars were separated by approximately one screen width, only one bar was visible at a time, producing 10 sequential bar passages. Following a 20-s inter-sweep blank interval, the same bar train traversed the display from right to left, producing 10 reverse-direction bar passages. **f.** Schematic of the VGA output used to track stimulus timing. Only pin 1 (red: red channel) and pin 6 (black: red channel ground) were used to create a custom BNC output, which was recorded continuously, providing an indirect measure of stimulus onset and offset timing. **g.** Expanded view of the first left-to-right bar passage within the forward sweep. Red dashed lines indicate the times at which the bar entered and left the visible display, corresponding to the change in the VGA red-channel signal. Camera-trigger pulses were generated continuously throughout the active unidirectional sweep. **h.** Expanded view of the final right-to-left bar passage within the reverse sweep. The red dashed line indicates the time at which the final bar left the display. The measured discrepancy between the final camera trigger and stimulus offset was 0.001 s—less than 0.0004% of the approximately 250-s validation recording—demonstrating stable synchronization between visual stimulus generation and camera acquisition. **i.** Schematic illustrating the appearance and disappearance of the checkerboard-patterned bar, aligned with the red dashed lines shown in panel g.

### Software Architecture and Firmware Implementation

ABISS is controlled by firmware developed in the Arduino IDE that we organized as two independent programs – each one optimized for auditory and visual intrinsic optical signal imaging respectively. Both firmware implementations use the same hardware platform while providing modality-specific stimulus generation and synchronized camera triggering. This approach allows the hardware to be rapidly reconfigured for different experimental paradigms by uploading the appropriate firmware without requiring hardware modification.

The auditory firmware uses Protothreads to interleave camera-trigger generation and tone-sequence control within a single non-preemptive loop, while a digital input is monitored in the main loop to start or stop the stimulation sequence. Timing in the auditory firmware is controlled using the Arduino micros() clock. One Protothread generates camera-trigger pulses at 20 Hz, while a second Protothread controls the tone sequence using microsecond-resolution timing targets. Because both image acquisition and stimulus reference the same microsecond clock timing source, stimuli remain precisely synchronized with camera acquisition throughout each recording session.

The visual firmware uses hardware timer interrupts to generate VGA synchronization signals while simultaneously rendering stimulus patterns and controlling camera triggering. Hardware timers generate horizontal and vertical synchronization signals, while RGB output is produced directly through GPIO pins. The display buffer represents a 440 × 480 pixel VGA-compatible output, with each byte corresponding to 8 horizontal pixels and each buffer row corresponding to 16 scanlines; this block-based representation reduces memory usage while still allowing high-contrast patterns, such as sweeping checkerboard bars, to be generated on an 8-bit microcontroller. The CPU enters idle sleep mode between interrupts to reduce unnecessary activity and preserve timing stability. Camera-trigger pulses are synchronized to the VGA refresh cycle by issuing one trigger pulse every third display frame, resulting in a 20 Hz acquisition rate from a 60 Hz display signal. Stimulus motion is updated at the same 20 Hz rate so that image acquisition and bar motion remain frame-locked throughout the experiment.

Experimental parameters are defined within the firmware and can be modified by editing user-accessible constants before uploading the program to the Arduino Nano. Auditory parameters include tone frequency, amplitude, pre-stimulus delay, stimulus duration, post-stimulus acquisition period, ramp step count and interval, inter-trial interval, and camera-trigger rate. Visual parameters include stimulus geometry, inter-stimuli spacing, stimulus speed, number of stimuli per sweep, sweep direction, inter-sweep interval, and the pattern rendered within each bar. The parameter values used in the present study are reported in the corresponding experimental Methods sections.

### Data Processing

Image processing and quantitative analyses were performed offline using custom MATLAB analysis scripts. Separate analysis pipelines were implemented for auditory and visual intrinsic optical signal imaging experiments to quantify stimulus-evoked cortical responses and generate functional maps.

### Auditory Intrinsic Signal Analysis

The acquired auditory IOSI data were systematically processed to compare the functional imaging map results from ABISS to those produced by the TDT RZ6. First, raw image files were sorted and segregated based on the stimulus delivery device.

For each tested tone frequency, including 3 kHz, 10 kHz, and 30 kHz, single-trial response maps were computed as the relative change in reflectance, Δ*R/R*. For each trial, baseline images were generated by averaging the first 30 frames (1.5 s), and response images were generated by averaging frames 41–80, corresponding to the post-stimulus response period. The relative reflectance change was calculated as:

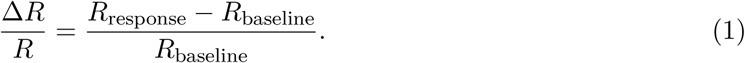

Each response map was spatially smoothed using a Gaussian filter (*σ*=3). For each device and frequency, response maps were averaged across trials to obtain the final cortical activation maps. Images displayed in the figures were contrast-normalized by clipping intensities to the 5th–95th percentiles.

To compare cortical activation patterns between ABISS and the commercial system, averaged response maps were converted into binary activation maps using a fixed thresholding procedure.

Response maps were first z-score normalized using the global image mean and standard deviation. Because intrinsic signal activation appears as a decrease in reflectance, activated pixels were defined as pixels below a threshold equal to 33% of the median of the lowest 25 z-scored pixel values. Similarity analysis was restricted to manually defined auditory responsive cortical regions.

Similarity between binary activation maps was quantified using the structural similarity index measure (SSIM). Cross-platform comparisons were performed between ABISS- and RZ6-generated binary maps for each stimulus frequency. To establish a commercial-system self-consistency reference for comparison, a separate RZ6-only dataset was divided into odd- and even-trial subsets, and SSIM was calculated between the corresponding binary response maps.

Bootstrap resampling was used to estimate the variability in SSIM measurements. For each stimulus frequency, 1000 bootstrap iterations were performed by randomly sampling 10 trials with replacement from the available trials for each condition. Averaged response maps, binary activation maps, and SSIM values were recomputed for every bootstrap iteration. Because the cross-system comparison and commercial self-consistency benchmark were obtained from different animals and recording sessions, no formal hypothesis testing was performed.

### Visual Intrinsic Signal Analysis

Retinotopic maps were generated using a standard phase-encoded Fourier analysis^14,20^. A cortical region of interest that spanned all visual areas but smaller than the entire field of view was manually defined from a reference image, and all subsequent analysis was restricted to this region. Image sequences were acquired during nine bidirectional sweep cycles for each stimulus orientation, yielding nine unidirectional sweeps in each of the four cardinal directions: left-to-right, right-to-left, top-to-bottom, and bottom-to-top. Slow fluctuations in illumination were removed by subtracting the mean intensity of each image frame. Each recording was divided into complete stimulus cycles based on the sweep period and acquisition rate. For each pixel, the complex response at the stimulus frequency was estimated by Fourier demodulation and averaged across cycles to obtain phase and response amplitude.

To compensate for the hemodynamic delay inherent to intrinsic optical signals, responses from opposing sweep directions were paired within each repetition by subtracting their phases and dividing the differences by two. The corrected phases were converted into vertical (elevation) and horizontal (azimuth) retinotopic maps and averaged across repeated acquisitions. Visual field-sign (VFS) maps were generated from the averaged retinotopic maps following Gaussian smoothing and gradient estimation. The visual field-sign at each pixel was calculated as the sine of the angular difference between the local azimuth and elevation gradients, yielding values between -1 and +1. Reversals in field-sign were used to identify boundaries between adjacent visual cortical areas.

### Code and Data Availability

All Arduino firmware and supporting files for ABISS and the custom MATLAB scripts used for the intrinsic-signal analyses are available in the Extended Data.

## Results

### ABISS integrates stimulus generation and synchronization into a unified control platform

ABISS was developed to simplify the distributed experiment-control pathway used in many intrinsic optical signal imaging (IOSI) workflows. In conventional implementations, sensory stimulation, camera triggering, trial control, and acquisition coordination are often handled by separate computers, commercial processors, data-acquisition systems, external pulse generators, or laboratory-specific software. In contrast, ABISS integrates these independent elements within a single programmable platform. An external trigger (user-initiated button press) initiates the experimental sequence, ABISS generates the auditory or visual stimulus, and the same firmware-defined timing pathway produces synchronized camera-trigger pulses for image acquisition. Importantly, ABISS only replaces the stimulus-control and synchronization infrastructure allowing seamless integration into an optical imaging workflow that is compatible with a multitude of cameras, speakers, or display devices. We first performed an engineering evaluation to determine whether ABISS could provide reliable stimulus generation and synchronization and once validated, we determined if it could fully support functional cortical mapping in both auditory and visual intrinsic optical signal imaging paradigms.

### ABISS provides reliable stimulus generation and synchronization for intrinsic optical signal imaging

Reliable synchronization between sensory stimulation and image acquisition is essential for intrinsic optical signal imaging because stimulus-evoked hemodynamic responses must be measured relative to precisely timed sensory events. We therefore first evaluated the engineering performance of ABISS by characterizing auditory stimulus generation and quantifying synchronization between stimulus presentation and camera triggering across extended recording sessions (Figure 3a).

Spectrogram analysis confirmed accurate generation of all three tone frequencies in the auditory paradigm, with distinct spectral peaks at 3, 10, and 30 kHz corresponding to the intended stimulus frequencies (Figure 3b). Camera trigger timing was measured at the beginning and the end of a continuous auditory session lasting approximately 1000 s. The measured pre-stimulus trigger interval for the first tone presentation was 2.00045 s, matching the intended 2-s delay. For the final tone presentation, the measured interval was 2.09 s, reflecting a cumulative timing drift of approximately 0.09 s across the ∼16.5 min experiment which corresponded to less than 0.01% of the total session duration (Figure 3c–d).

We next evaluated synchronization during visual stimulation by simultaneously recording the camera-trigger signal and channel voltage output from a single channel (red) of the VGA during a standard length visual mapping session (Figure 3e-f). Voltage from the red channel increased as the checkerboard-patterned stimulus entered onto the display and returned to baseline as the stimulus left the screen, providing an indirect measure of visual stimulus timing (Figure 3g, i). By the final trial, the discrepancy between stimulus offset and the end of the camera trigger was 0.001 s across the approximate 250 s duration of a standard experiment, corresponding to less than 0.0004% timing error associated with drift (Figure 3h). Together, these measurements demonstrate that ABISS maintains reliable synchronization between stimulus delivery and camera triggering that is well aligned for measuring seconds-long hemodynamic responses throughout both auditory and visual imaging paradigms.

### ABISS reliably supports acquisition of auditory functional imaging maps

Having verified accurate stimulus generation and precise synchronization with image acquisition, we next asked whether ABISS could support routine auditory cortical mapping. Auditory IOSI presents an additional practical challenge because mapping in animal models often requires calibrated high-frequency or ultrasonic tone generation beyond the reliable output range of typical consumer-grade computer audio hardware. As such, studies generally rely on specialized sound-generation hardware and laboratory-specific control software, which provide high output fidelity but increase cost and system complexity^3,16,21^. To determine whether ABISS could provide the auditory signal generation and stimulus control needed for routine auditory cortical mapping, we directly compared auditory intrinsic signal maps evoked by ABISS-generated tones with those evoked by a commercial TDT RZ6 system.

Because intrinsic signal maps vary across animals and recording sessions, we designed two complementary validation experiments. In the first experiment, tonotopic maps produced from ABISS outputs and those from the RZ6 system were compared directly within the same imaging session in the same animal under identical experimental conditions. Both systems were calibrated to deliver 75 dB SPL stimuli before testing, and trials alternated between the two stimulation systems while maintaining the same speaker, acoustic delivery path, imaging setup, and animal preparation. In the second experiment, we compared maps generated from odd and even trials of the RZ6 system to establish a commercial-system self-consistency benchmark. This would allow us to capture differences in functional maps that could only be explained by trial-by-trial variability in optical responses.

The response maps shown in Figure 4a–g were all obtained from the direct ABISS–RZ6 comparison experiment. The composite tonotopic map demonstrates orderly frequency-dependent activation across the tested frequencies (Figure 4a). ABISS-generated responses at 3, 10, and 30 kHz produced spatially organized activation patterns within the auditory cortex (Figure 4b–d). Corresponding RZ6-evoked responses exhibited similar activation locations and spatial organization (Figure 4e–g), indicating that ABISS-generated stimuli can produce highly consistent frequency-dependent intrinsic responses comparable to those generated with a dedicated commercial system.

**Figure 4.**
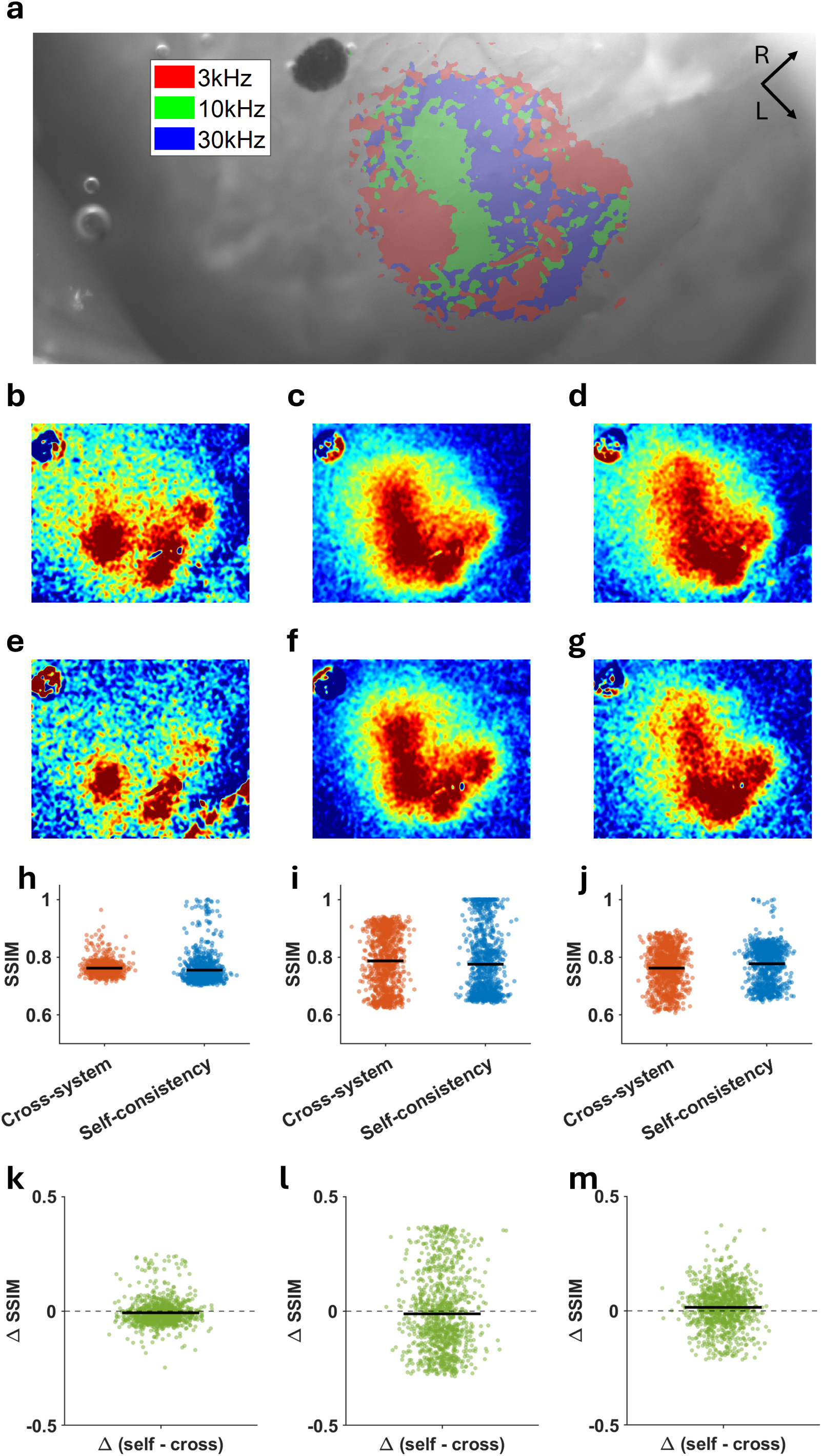
ABISS-generated auditory stimulation produces intrinsic signal maps comparable to a commercial TDT RZ6 system. **a.** Composite auditory intrinsic signal map overlaid on the anatomical reference image. Responses to 3, 10, and 30 kHz tones are shown in red, green, and blue, respectively. **b–d.** ABISS-evoked intrinsic response maps for 3 kHz (b), 10 kHz (c), and 30 kHz (d) tones independently. **e–g.** RZ6-evoked intrinsic response maps for 3 kHz (e), 10 kHz (f), and 30 kHz (g) tones independently. **h–j.** Bootstrap distributions of SSIM values comparing ABISS–RZ6 cross-system similarity and RZ6 odd–even self-consistency for 3 kHz (h), 10 kHz (i), and 30 kHz (j). Each point represents one bootstrap resample; black bars indicate the mean SSIM. **k–m.** Bootstrap distributions of ΔSSIM, defined as SSIM_RZ6 SELF_ − SSIM_ABISS_*_−_*_RZ6_, for 3 kHz (k), 10 kHz (l), and 30 kHz (m). The dashed horizontal line indicates zero difference between cross-system similarity and RZ6 self-consistency.

To quantify map similarity, response maps were converted to binary activation maps and compared using structural similarity index measure (SSIM). For each frequency, 1000 bootstrap samples were generated by resampling trials with replacement, and SSIM values were calculated for both the ABISS-RZ6 comparison and the RZ6 self-consistency benchmark. Across all tested frequencies, ABISS–RZ6 similarity closely matched the RZ6 self-consistency benchmark (Figure 4h–j). Median ΔSSIM values were -0.012 for 3 kHz, -0.036 for 10 kHz, and 0.017 for 30 kHz (Figure 4k–m), and all 95% bootstrap intervals included zero. Thus, the differences between ABISS-generated and RZ6 generated maps were no larger than the variability observed between trials comparing odd and even trials from the commercial system. Together, these findings demonstrate that ABISS combines high resolution auditory stimulus generation and trial synchronization that is capable of producing IOSI functional maps comparable to established commercial stimulation workflows.

### ABISS supports visual intrinsic optical signal imaging

After establishing that ABISS supports auditory intrinsic optical signal imaging, we next asked whether the same hardware architecture could generalize to visual cortical mapping. Specifically, we evaluated whether the integrated VGA stimulus-generation and synchronized camera-triggering pathways implemented in ABISS could generate visual stimuli that would be essential for standard retinotopic imaging. Prior visual IOSI studies have commonly relied on computer-based visual stimulus software, including Psykinematix, MATLAB/Psychophysics Toolbox, PsychoPy, or custom stimulus programs, while image acquisition and synchronization are controlled through separate workflows (camera software, acquisition computers, data-acquisition hardware, or external trigger interfaces) ^13,14,16,18,19,22^. Although these approaches are powerful and flexible, they require synchronization across multiple stimulation and acquisition components. We therefore asked whether ABISS could simplify this workflow for basic retinotopic mapping by generating VGA-based drifting checkerboard-bar stimuli and synchronized camera-trigger pulses from the same Arduino-controlled board.

Horizontal and vertical retinotopic maps were generated using phase-encoded Fourier analysis of intrinsic signal responses evoked by drifting checkerboard bars moving in opposite directions. The horizontal retinotopic map showed a smooth progression across the imaged cortical region, indicating spatially organized azimuth representation (Figure 5a). The corresponding vertical retinotopic map similarly revealed structured elevation-dependent responses (Figure 5b). These maps were subsequently combined to generate a visual field-sign map based on the angular relationship between local horizontal and vertical retinotopic gradients. The resulting visual field-sign (VFS) map showed alternating mirror and non-mirror representations characteristic of the mouse visual cortex (Figure 5c).

**Figure 5.**
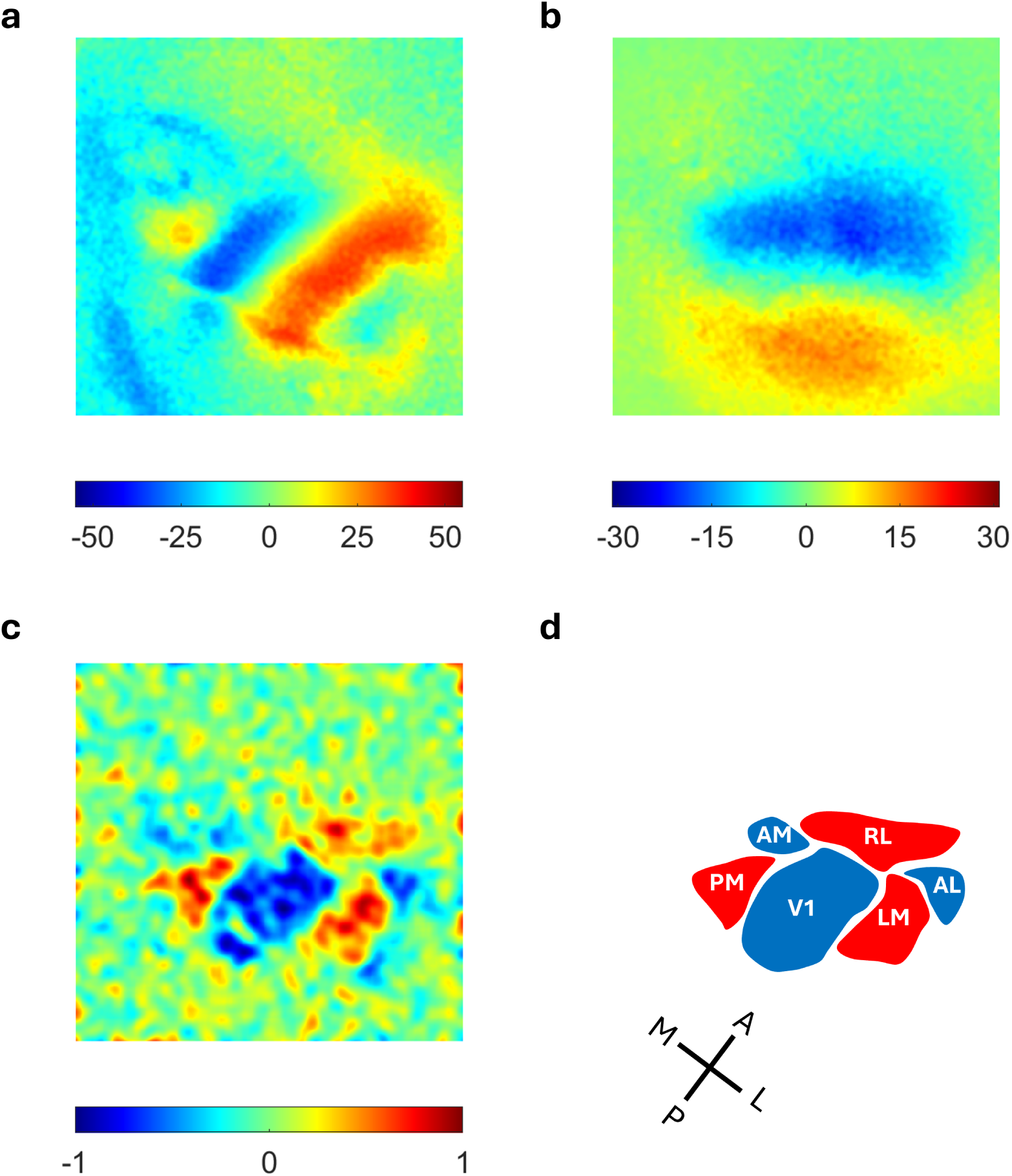
ABISS-driven visual stimulation supports retinotopic mapping of the mouse visual cortex. **a.** Horizontal retinotopic map generated from phase-encoded intrinsic signal responses to drifting-bar visual stimulation. Color code indicates estimated azimuth position in visual degrees. **b.** Vertical retinotopic map generated from responses to drifting bars moving along the vertical axis. Color code indicates estimated elevation position in visual degrees. **c.** Visual field sign map computed from the local gradients of the horizontal and vertical retinotopic maps. Positive and negative values indicate opposite visual field representations. **d.** Manual delineation of putative visual cortical areas based on field-sign reversals and comparison with previously reported mouse visual cortex maps from IOSI studies ^1,14,18^. Mirror-image representations are shown in blue, and nonmirror-image representations are shown in red. Abbreviations: V1, primary visual cortex; LM, lateromedial area; AL, anterolateral area; RL, rostrolateral area; AM, anteromedial area; PM, posteromedial area.

Based on these field-sign reversals and the expected spatial arrangement of mouse visual areas described in previous IOSI studies ^1,14,18,23^, we identified putative V1 and higher visual cortical areas, including LM, AL, RL, AM, and PM (Figure 5d) from the VFS in Figure 5c. Mirror-image representations were labelled in blue and nonmirror-image representations in red. The recovery of expected retinotopic gradients and visual field-sign organization demonstrates that ABISS supports visual stimulation generation conducive to functional mapping of mouse visual cortex.

### ABISS supports routine functional mapping across sensory modalities

Collectively, these experiments establish that ABISS provides an integrated multimodal control platform well-suited for basic intrinsic optical signal imaging. Engineering validation demonstrated precision in the timing of auditory and visual stimulus generation together with reliable synchronization between stimulus delivery and image triggering throughout extended recording sessions. Biological validation further demonstrated that these engineering characteristics translated into the production of functional cortical maps comparable to those produced by established workflows in the auditory and visual domains. Together, these findings demonstrate that ABISS provides the stimulus control, and timing precision to support routine intrinsic optical signal imaging across both auditory and visual sensory modalities.

## Discussion

We developed and validated ABISS, an open-source hardware and software platform that reduces the practical barriers to routine functional cortical mapping using intrinsic optical signal imaging. Although ABISS could be used for traditional IOSI, it was not designed to replace high-end systems in experiments where IOSI is itself the primary biological measurement. Instead, it is intended for a common use case in systems neuroscience, where IOSI could serve as a critical early step to localize functional cortical areas before downstream experiments that require accurate targeting of specific regions. In this role, IOSI helps the experimenter move beyond stereotaxic coordinates alone by enhancing identification of regional boundaries of interest in the same animal that will later undergo electrophysiology, viral injection, pharmacology, lesioning, or other circuit-level manipulations. For this purpose, the most important requirement is not maximal stimulus complexity or imaging performance, but a reliable, reproducible, and easily deployed method for mapping functional regions in individual animals. By consolidating stimulus generation, trial timing, and acquisition triggering into a low-cost and portable hardware system, ABISS makes functional mapping substantially easier to adopt across laboratories that will also provide benefit by increasing reproducibility in systems neuroscience.

Several modern imaging techniques can provide functional cortical activity mapping, including calcium imaging^24,25^, functional magnetic resonance imaging (fMRI)^26,27^, and functional ultrasound localization imaging ^28^. These methods provide powerful capabilities, including cellular or genetically defined activity measurements, non-invasive whole-brain hemodynamic mapping, and sensitive vascular imaging. However, they often require specialized instrumentation, use of viral vectors or reliance on transgenic lines, magnetic resonance imaging infrastructure, or dedicated ultrasound acquisition hardware and analysis workflows. Such requirements may be disproportionate when the immediate objective is to simply identify a functional cortical region before subsequent electrophysiological recording or another targeted experiment where precision is essential. For this preparatory application, intrinsic optical signal imaging occupies a particularly useful niche: it enables rapid, minimally invasive, wide-field functional localization using relatively simple optical instrumentation that does not rely on exogenous dyes or genetically encoded reporters ^4–6^. Recent work emphasizes that biological variability in cortical geography limits the precision that can be expected using atlas-based targeting ^3^. Consequently, the primary obstacle to routine IOSI is often not the imaging methodology itself but the practical complexity of coordinating stimulus generation, synchronization, and experimental control. ABISS was developed specifically to reduce this barrier to entry while preserving the simplicity and flexibility that have made intrinsic optical signal imaging a widely adopted method for functional cortical mapping.

The engineering characterization demonstrates that ABISS provides the temporal precision necessary for intrinsic optical signal imaging. Across extended recording sessions, stimulus generation remained spectrally accurate and synchronization between stimulus delivery and camera triggering remained consistent. Although the absolute timing accuracy of modern commercial processors exceeds the requirements of most IOSI experiments, intrinsic imaging primarily depends on maintaining reliable temporal relationships between stimulus presentation and image acquisition throughout long recording sessions. The observed stability of ABISS therefore indicates that a microcontroller-based platform can reliably support intrinsic imaging protocols without requiring more sophisticated or costly synchronization hardware.

Having established ABISS as capable of high precision stimulus generation and timing control, we further validated ABISS as a practical functional-localization tool. In the auditory cortex, intrinsic signal maps generated using ABISS were spatially comparable to maps generated using a commercial auditory system, with structural similarity values consistent with those produced from alternating trials using the commercial system alone. Importantly, the goal of this comparison was not to demonstrate that ABISS reproduces every capability of a commercial auditory processor, but rather to determine whether the platform provides stimulus control and auditory stimulus capability to produce reliable functional localization. We further demonstrated that the same hardware platform supports visual retinotopic mapping, generating retinotopic gradients and visual field-sign organization consistent with the expected organization of the mouse visual cortex ^18,23^. Together, these experiments show that ABISS generalizes across sensory modalities while preserving the timing sensitivity required for multimodal functional cortical mapping.

Recent efforts have substantially improved the accessibility of IOSI through inexpensive illumination systems, open-source analysis software, and simplified optical hardware^18,19^. These efforts have simplified IOSI adoption, but many implementations still rely on separate systems for stimulus generation, image acquisition, and synchronization. ABISS was designed to address a complementary aspect of the workflow by integrating several commonly separated experimental-control functions into a single reproducible hardware pathway. Many IOSI workflows depend on commercial systems, external synchronization hardware, or laboratory-specific software to generate stimuli and coordinate trial timing. Commercial systems remain advantageous when experiments require sophisticated visual rendering, multichannel auditory stimulation, closed-loop behavioral control, or integration with large-scale acquisition systems. In contrast, ABISS was developed for laboratories whose primary objective is to reliably identify functional anatomical boundaries before downstream experiments. To our knowledge, ABISS is the first low-cost, single-device platform capable of integrating auditory stimulus generation, visual stimulus generation, trial timing, and camera-acquisition triggering for IOSI. This distinction defines the experimental situations for which ABISS is most appropriate. The platform is particularly well suited for routine auditory or visual cortical mapping, teaching laboratories, shared imaging facilities, and research groups seeking to implement intrinsic optical signal imaging without purchasing specialized commercial control hardware. Because experimental protocols are defined in firmware, investigators can readily modify stimulus timing, auditory frequencies, visual stimulus geometry, and trial structure without redesigning the underlying hardware. These characteristics make ABISS especially useful where simplicity, portability, and reproducibility are more important than maximal stimulus complexity.

ABISS is also inexpensive and simple to modify. The current design can be assembled from commonly available components for less than $150.00 USD, excluding external experimental equipment such as the camera, illumination source, video stimulus monitor, and speaker. Experimental protocols can be modified by editing the Arduino firmware and uploading new code, allowing users to adjust stimulus frequency, amplitude, timing, visual pattern structure, camera-triggering rate, and trial organization without building a separate computer-control interface. The device is powered by direct current and contains no moving components such as cooling fans, making it well suited for acoustically sensitive environments, including sound-attenuating booths or other settings where hardware noise is undesirable. Its small physical footprint (4 × 3 × 1 inches), and low weight (150 g), allow it to be transported between laboratory spaces and deployed in settings where moving large equipment is impractical, such as vivarium’s, shared surgical suites, or experimental procedure rooms. Together, these features make ABISS easy to deploy and readily usable as a dedicated portable system for routine functional mapping.

The limitations of ABISS should be interpreted in light of the intended role of the device. The Arduino Nano provides sufficient computational performance for the timing requirements of intrinsic optical signal imaging but is not intended for computationally intensive stimulus generation. The current VGA output is appropriate for high-contrast retinotopic stimuli but is insufficient to produce naturalistic visual scenes, high-resolution graphics, or environments where dynamically corrected display is required. Similarly, the present auditory implementation supports single-channel output and therefore is not currently capable of binaural stimulation or complex spatial auditory experiments. These limitations reflect deliberate design decisions to maximize the capabilities of the Arduino microprocessor rather than technical shortcomings. The objective of ABISS is not to replace specialized commercial systems for demanding sensory neuroscience experiments but to provide a practical and reproducible platform for routine functional localization.

Future development of ABISS could substantially expand the range of experimental applications while preserving its open-source architecture. More capable microcontrollers could support higher-resolution visual stimuli, increased computational flexibility, additional channels, and more sophisticated auditory paradigms. Integration of graphical interface software, automated calibration routines, and network-based communication would further simplify adoption by laboratories with limited engineering experience. Equally important will be broader validation across additional sensory systems, animal species, and independent laboratories, providing a more comprehensive assessment of reproducibility across diverse experimental environments.

The broader significance of ABISS extends beyond the development of a new stimulation device. As biological variability across animals limits the precision of atlas-based targeting, routine per-animal functional mapping is becoming a more important component of systems neuroscience workflows. Widespread adoption of intrinsic optical signal imaging has historically been constrained less by the optical instrumentation than by the complexity of integrating stimulation, synchronization, and experimental control into a simple and easy-to-deploy workflow. By consolidating these functions into a compact, inexpensive, and openly available platform, ABISS lowers the practical barrier to implementing routine functional cortical mapping. We anticipate that reducing these barriers will broaden access to IOSI, improve the reproducibility of targeted neuroscience experiments across laboratories, and encourage routine per-animal functional mapping before downstream electrophysiological and circuit-manipulation studies in which anatomical precision is critical.

## Code and Data Availability

All Arduino firmware and supporting files for ABISS are publicly available at https://github.com/TylerMarr/ABISS. The custom MATLAB scripts used for the intrinsic-signal analyses are available at https://github.com/Zhili-Qu/ABISS-IOSI-analysis, and the imaging datasets are available from the corresponding author upon reasonable request.

## Author Contributions

Z.Q. performed all imaging experiments and data analysis. Z.Q., K.K., T.W., and S.N.D. contributed to the overall hardware design. K.K. designed the printed circuit board. T.W. and S.N.D. contributed to the design and selection of electronic components. T.W. and T.A.M. wrote the software. Z.Q. and H.J.G. wrote the manuscript. M.G. provided supervision and advice. H.J.G. conceived the study, supervised the project, and acquired funding.

## Acknowledgment

We thank Bo Yuan for designing the custom enclosure for ABISS. This research was supported by National Science Foundation grants SMA-2319321 (H.J.G. and K.S.), EFRI-2515342 (H.J.G. and M.G.), and NSF Expedition in Computing “Mind in Vitro” (IIS-2123781; M.G.), as well as Brain Research Foundation Award BRFSG-2023-13 (H.J.G.).

## Declaration of AI in Scientific Writing

During the preparation of this work, the authors used Claude (Anthropic) and ChatGPT (OpenAI) to improve the readability and language of the manuscript. After using these tools, the authors reviewed and edited the content as needed and take full responsibility for the content of the publication.

## Extended Data

**Extended Data Table 1:**
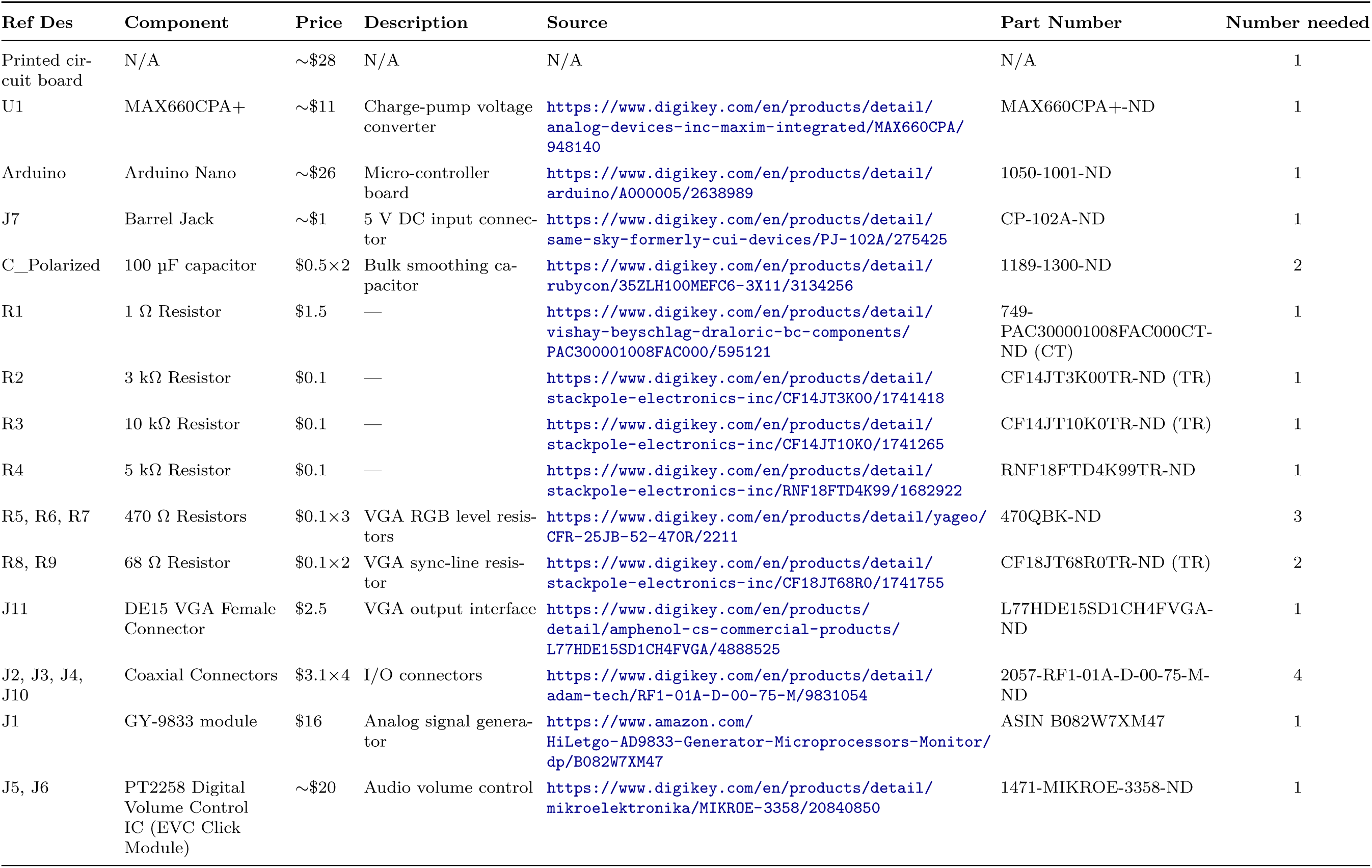
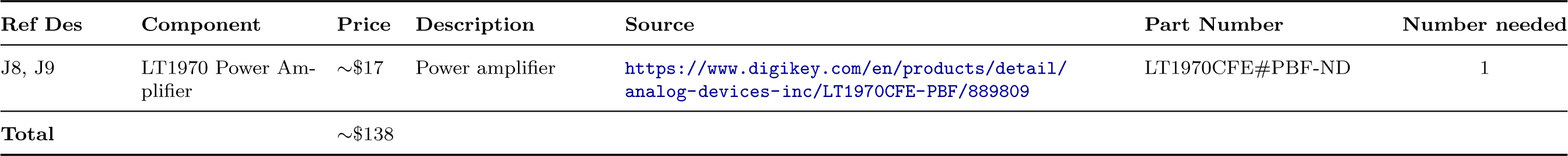
Assembly and bill of materials. Prices are approximate and exclude external experimental equipment (camera, illumination source, monitor, speaker). Source links point to the exact part listing used.

**Extended Data Figure 1-1.**
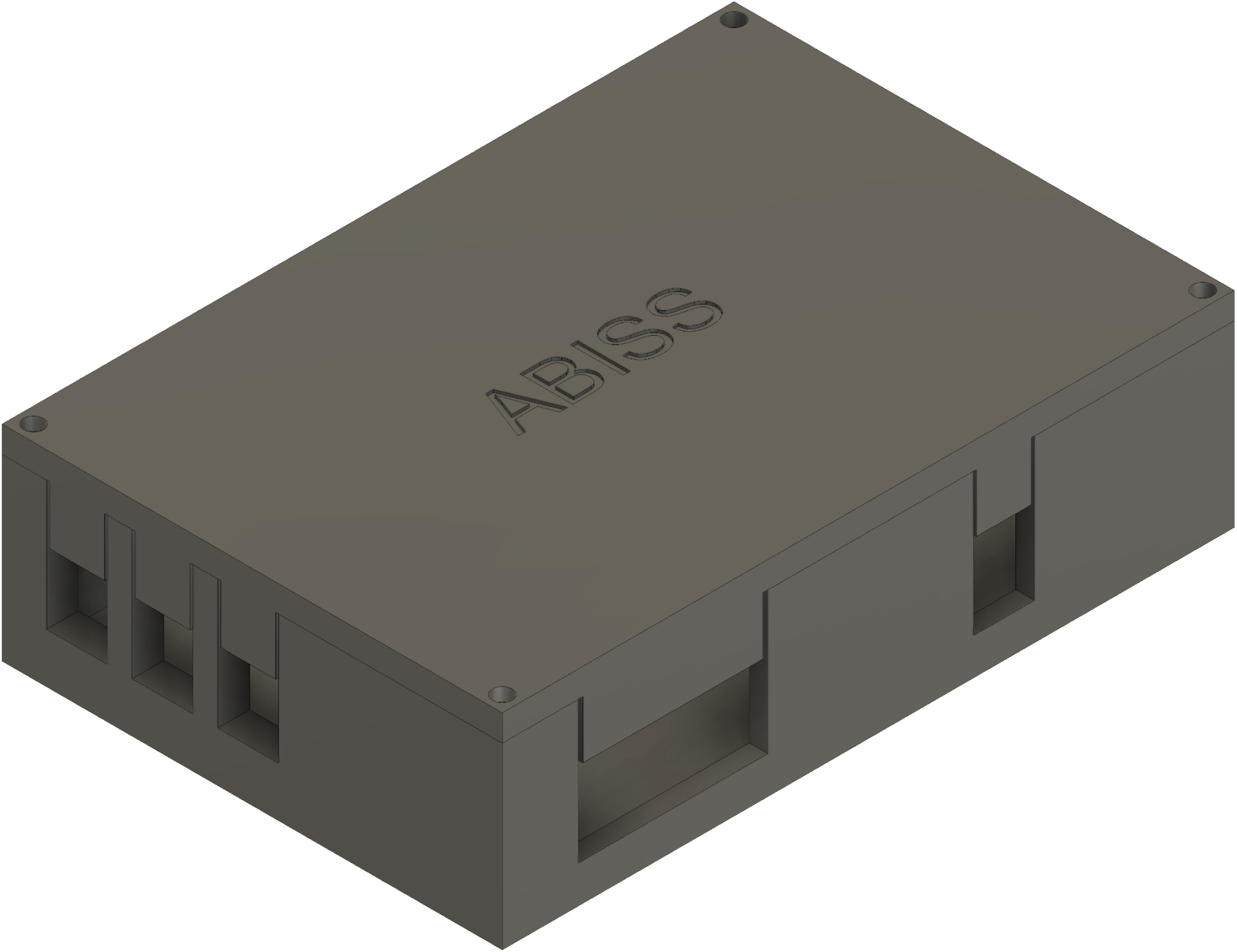
CAD schematic of the custom enclosure designed for ABISS. The enclosure protects the board from dust and debris while remaining fully functional with window openings that expose all input/output connectors for external access. Four corner screws secure the lid, allowing the enclosure to be opened or closed as needed to access or remove the ABISS board.

**Extended Data Figure 1-2.**
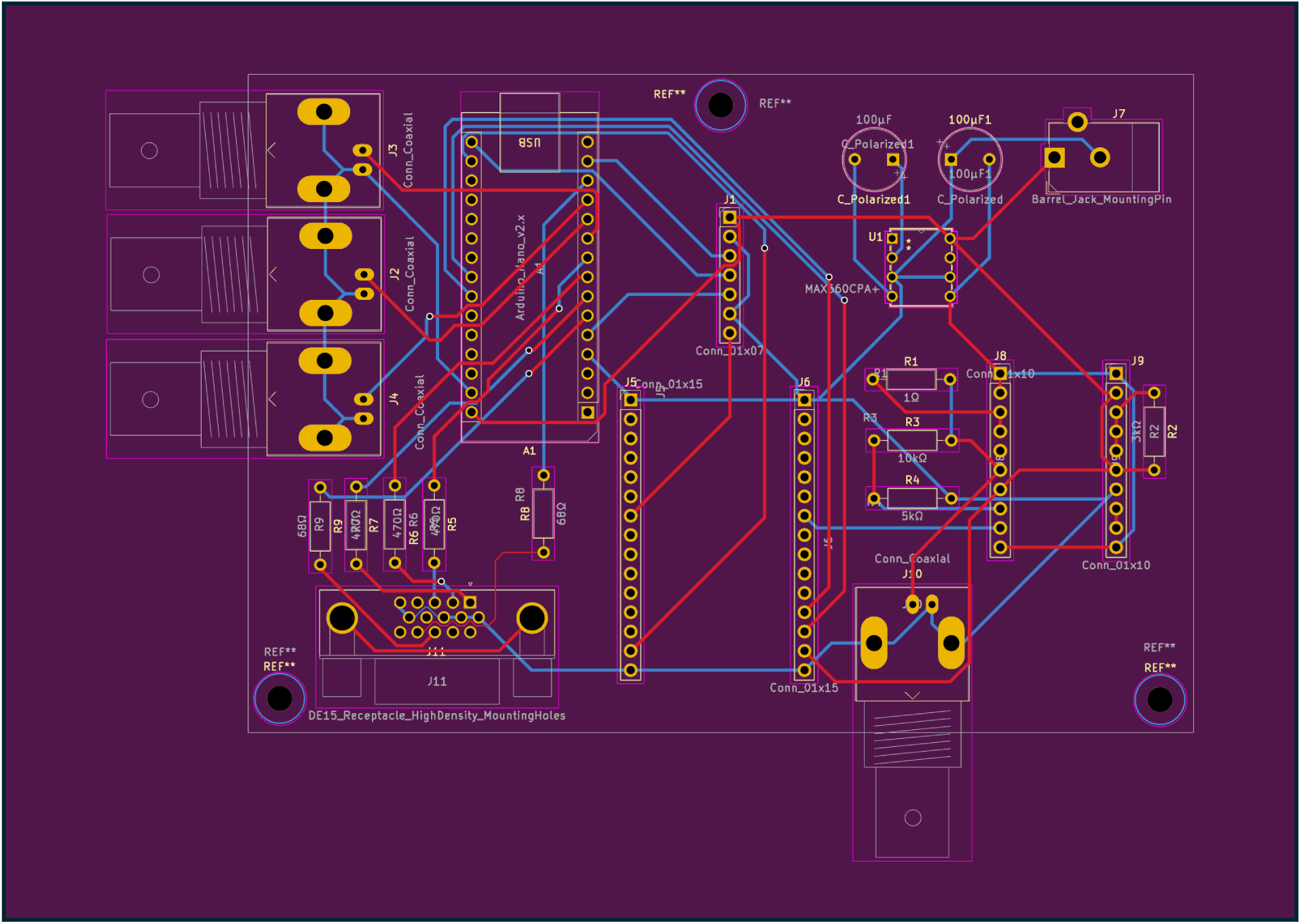
Customized PCB layout. PCB architecture showing component placement, through-hole footprints, and routed copper traces for the ABISS board.

**Extended Data Figure 1-3.**
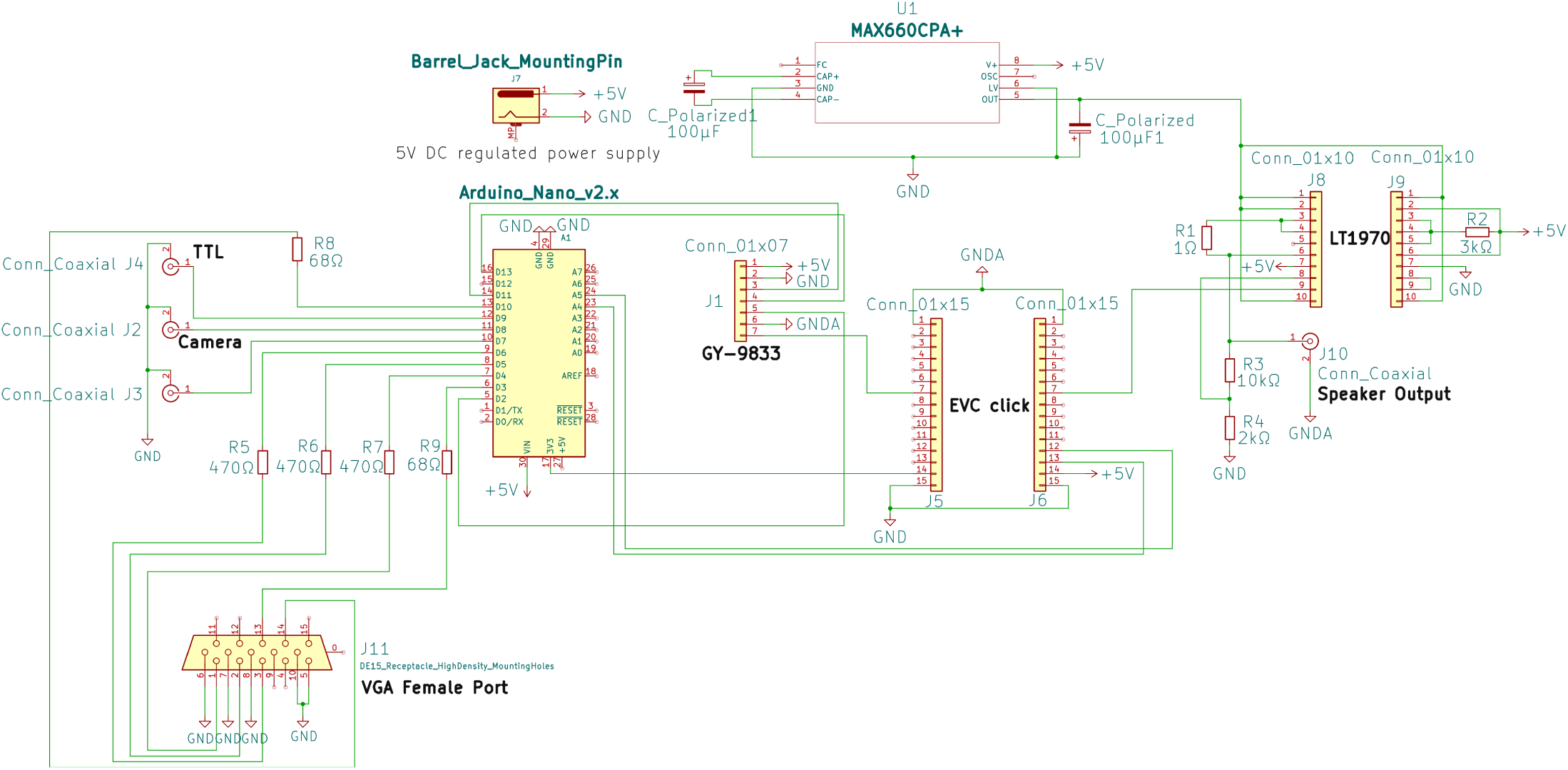
Complete circuit schematic of the ABISS board. Full KiCad-generated schematic showing every component and connection in the ABISS system.

